# Quantitative live cell imaging of nuclear shape and chromatin dynamics during development and environmental stress in *Arabidopsis thaliana*

**DOI:** 10.64898/2026.02.27.708654

**Authors:** Joh Demura-Devore, M. Arif Ashraf

**Affiliations:** Department of Botany, University of British Columbia, Vancouver, BC, Canada

## Abstract

The nucleus is the characteristic organelle for eukaryotic organisms. Unlike the classic textbook view of static two-dimensional nuclei, nuclear shape is dynamic inside the live cell. The alteration or deformed nuclear shape is the hallmark of cancer in animal cells and environmental stress in plants. The nuclear envelope proteins interact with chromatin to regulate gene expression. Unfortunately, we have limited knowledge about the impact of abiotic stress on nuclear shape, movement, and chromatin dynamics. To circumvent this issue, we are utilizing a dual fluorescently tagged marker lines – nuclear envelope protein and chromatin – to perform live cell imaging in the model plant *Arabidopsis thaliana* root. The live cell imaging was performed in control and salt-stressed conditions. We utilized these captured movies to analyze through open-source image processing software Fiji/ImageJ with the help of the TrackMate plugin. Using this method, we have demonstrated that chromatin velocity is decreased in salt-treated conditions. This method will be widely applied to quantitative live cell imaging of nuclear shape and chromatin dynamics during plant development and environmental stress.

**Summary:** This process aims to simultaneously record nucleus and chromatin dynamics in *Arabidopsis thaliana* roots and investigate changes in these dynamics in response to developmental and environmental cues.

## Introduction

Nuclear shape is variable between different cell types and during the course of development in *Arabidopsis thaliana* roots (1,2). For instance, nuclear shape is spherical in the actively dividing meristematic regions. In contrast to the meristematic region, the nucleus in the transition and elongation zone is less spherical. Furthermore, nuclei in differentiated cells such as root hairs pass through the tight space and are actively transformed into elongated nuclei (3,4). The dynamic changes in nuclear shape and movement are crucial to the functioning of the cell (5,6). Defects in nuclear shape lead to functional impairment such as reduced nuclear movement (3,7). In mammalian cells, changes in cell and nucleus shape are directly correlated with changes in gene expression and protein synthesis (8).

During interphase, a cell’s DNA is organized as chromatin. The DNA in chromatin is bundled around histones, which are able to tighten and loosen the DNA to alter expression. Similarly, chromatin positioning relative to the nuclear envelope and proximity to the nuclear periphery are also important for gene expression (9). Environmental stress, such as osmotic stress, alters nuclear shape and consequently changes gene expression in a set of touch-sensitive gene expression in *A. thaliana* roots (10). In this context, studying the nuclear shape and chromatin dynamics through live cell timelapse imaging is important.

In this method, we are presenting a live cell imaging technique to visualize the nuclear shape (fluorescently tagged outer nuclear envelope protein WIP1) and chromatin (cenH3) dynamics simultaneously. Previously, nuclear shape and chromatin dynamics were imaged and quantified separately; as a result the relationship between nuclear shape and chromatin dynamics remained unexplored. Additionally, we studied the chromatin dynamics in a quantitative manner by using the TrackMate plugin of open-source ImageJ/Fiji software. We tested a dual fluorescent line and chromatin dynamics techniques presented here under salt stress conditions and discovered an alteration in chromatin morphology and dynamics due to stress conditions. We believe our method will be widely applicable beyond plant model systems, because chromatin and the nuclear envelope are present widely across eukaryotes in the tree of life.

1. **Growing of *A. thaliana* Seedlings**
  1.1. Growth Media Preparation
    1.1.1. Create ½ MS 1% sucrose 1% agar growth media.
    1.1.2. Create ½ MS 1% sucrose 1% agar 100mM NaCl growth media.
    1.1.3. While autoclaving growth media, also autoclave water.
    1.1.4. Growth media can be stored in a 4°C fridge and microwaved to liquify.
  1.2. Setting Growth Plates and Seeds
    1.2.1. In a flow hood, use a pipette gun to pipette 50mL of growth media into each of ten plates (**Figure 1A**).
    1.2.2. While the media solidifies, clean WIP1-GFP x CENH3-mRFP seeds by flooding them in an Eppendorf tube with 1mL 70% ethanol for 10 minutes (**Figure 1B**).
    1.2.3. After 10 minutes, in the flow hood, remove the ethanol and rinse the seeds twice with 1mL autoclaved water (**Figure 1C**).
    1.2.4. Once the media has set, use a 1000μL pipette to place cleaned seeds onto growth plates in two rows of 14 seeds per plate (**Figure 1D**).
    1.2.5. Label the plates with sharpie and seal them with Micropore tape before leaving the flow hood (**Figure 1E**).
  1.3. Germinating Seedlings
    1.3.1. Wrap growth plates in aluminum foil and cold stratify them in a 4°C fridge for a minimum of 2 days (**Figure 1F**).
    1.3.2. After cold stratification, remove plates from the fridge and leave them vertically under growth lights for a minimum of 4 days (**Figure 1G**).
    1.3.3. Transfer half of plates to 100mM NaCl growth media in flow hood, seal and label plates (**Figure 1H**).
    1.3.4. Grow plates under grow lights for 2 days before imaging (**Figure 1I**).
2. **Live Cell Imaging of *A. thaliana* Root Nuclei and Chromatin**
  2.1. Slide Preparation
    2.1.1. Pipette 60μL MQ water onto a glass slide (**Figure 2A**).
    2.1.2. Lay 3 *A. thaliana* seedlings’ roots into the water (**Figure 2B**).
    2.1.3. Place a 22x22mm #1.5 slide cover over the roots without covering the leaves (**Figure 2C**).
  2.2. Confocal Microscope Settings
    2.2.1. Set laser exposure time to 1000ms (1 second).
    2.2.2. In your laser combiner, set the 488nm laser to 50% for GFP and the 561nm to 50% for RFP (**Figure 2D**). If recording for longer than 10 minutes, you may decrease the intensity to reduce photobleaching.
    2.2.3. Set Z-slice step size to 0.3um.
    2.2.4. Set interval to 1 minute and total duration to 10 minutes.
  2.3. Imaging
    2.3.1. Locate roots and select ∼20 z-slices. Selecting more than 20 z-slices may prevent each set of images from being taken within 1 minute, but can be done if desired.
    2.3.2. Start timelapse.
    2.3.3. Move data to a computer for analysis.
3. **Tracking and Analysis of Data** Note: You will need the Fiji version of ImageJ, which comes with TrackMate preinstalled.
  3.1. Initialization
    3.1.1. Open ImageJ/Fiji, then click and drag your .vsi file onto the ImageJ/Fiji application window (**Figure 3A**).
    3.1.2. You will be shown a prompt titled “Bio-Formats Import Options”, click “Ok” in the bottom right (**Figure 3B**).
    3.1.3. You will be shown a prompt titled “Bio-Formats Series Options”. Make sure “Series 1” is selected and click “Ok” in the bottom left (**Figure 3C**). NOTE: The file may take a minute to load, especially if it has many Z-layers or time points.
    3.1.4. Once your image has opened (**Figure 3D**), split the signals by navigating to image, color and split (**Figure 3E**). This will create two separate windows (**Figure 3F**).
  3.2. Preprocessing Chromatin
    3.2.1. Click on your chromatin window (**Figure 3F**) and apply the Z-project operation by navigating to image, stacks and “Z project…” (**Figure 4A**). Change projection type to “Max Intensity” and make sure “All Time Frames” is active, then click “Ok” (**Figure 4B**). NOTE: This step allows for easier viewing, but if you intend to analyze Z-dimensional data, skip this step.
    3.2.2. For visibility, adjust the contrast by pressing Ctrl+Shift+C or navigating to image, adjust, and “Brightness/Contrast” (**Figure 4C**). Slide the second bar (labeled “Maximum”) to the left until the chromatin is easily visible (**Figure 4D**). Click apply, which will trigger a warning that pixel values will be adjusted, press “Ok”, then you will get a popup asking to apply the LUT to all stack slices, press “Yes” (**Figure 4E**).
  3.3. Tracking Chromatin
    3.3.1. Navigate to plugins, tracking, and TrackMate (**Figure 5A**). You may need to click the down arrow in the plugins menu to scroll to the tracking option (**Figure 5A**).
    3.3.2. You will be shown the initial settings. Confirm that the calibration settings are accurate, then click “Next” (**Figure 5B**).
    3.3.3. Select “Thresholding Detector” and click “Next” (**Figure 5C**).
    3.3.4. On your image window, move to the last frame of your timelapse (**Figure 5D**). Next to the intensity threshold, click “Auto”, then click “Preview”. Confirm that the preview includes your chromatin and doesn’t include debris or noise. If necessary, manually adjust the intensity threshold. Then, click “Next” (**Figure 5D**).
    3.3.5. Initial thresholding detection will occur, click “Next” (**Figure 5E**).
    3.3.6. If your data is noisy, manually threshold or click “Auto”. Then, click “Next” (**Figure 5F**).
    3.3.7. This page allows filtering of spots. Ignore this and click “Next” (**Figure 6A**).
    3.3.8. Select “Simple LAP tracker” and click “Next” (**Figure 6B**).
    3.3.9. Set linking max distance to 5 microns, gap-closing max distance to 15 microns, and gap-closing max frame gap to 1. Click “Next” (**Figure 6C**).
    3.3.10. Tracking will occur. You can view the tracks on the original image window. Click “Next” (**Figure 6D**).
    3.3.11. This page allows track filtering. If you have a lot of spots created by noise, press the green “+” button in the bottom left to create a filter, then choose the “Number of spots in track” mode and press “Auto” (**Figure 6E**). Make sure the mode is set to “Above”. This will eliminate spots that quickly appear and disappear.
    3.3.12. The display options page allows you to export your data as a csv. At the bottom of the page, click the “Tracks” button, then “Export to CSV” (**Figure 7A**).
    3.3.13. This page allows built-in plotting, ignore it and click “Next”.
    3.3.14. On the final page, you may save the video using “Capture overlay” and pressing “execute”. This will create a new image series that you can save by clicking on it, then navigating to file, save as, “GIF…” (**Figure 7B**).
  3.4. Basic Velocity Analysis
    3.4.1. Open your tracks CSV in an Excel-like software.
    3.4.2. Delete all columns except for TRACK_ID, POSITION_X, POSITION_Y, POSITION_T, FRAME, and RADIUS.
    3.4.3. The spots in each track will be out of order. To order them, first freeze the top four rows and then create a new column titled “TRACK_FRAME” and copy paste this formula into the column: =concatenate(A4, “_”, if(E4>9, E4, concatenate(“0”, E4))).
    3.4.4. Select the TRACK_FRAME column, right click it, then click “Sort sheet A-Z” or the equivalent in your software of choice. The spots within each track should now be out of order.
    3.4.5. To confirm no spots are missing in the middle of any tracks, create a new column and use this formula: =if(E4>E3, E4-E3, “NA”). The first spot of each track should return “NA” and the rest should be 1.
    3.4.6. Create a column for Velocity (micron/frame) and paste the following formula into it: =if(H4=“NA”, “NA”, sqrt((B4-B3)^2 + (C4-C3)^2)). This will use the pythagorean theorem to calculate the distance between two points.
    3.4.7. To translate this to Velocity (micron/second), create a new column and paste the following formula into it: =IF(I5=“NA”, “NA”, I5/(D5/E5)). This will find the seconds/frame and translate microns/frame accordingly.

**Figure 1.**
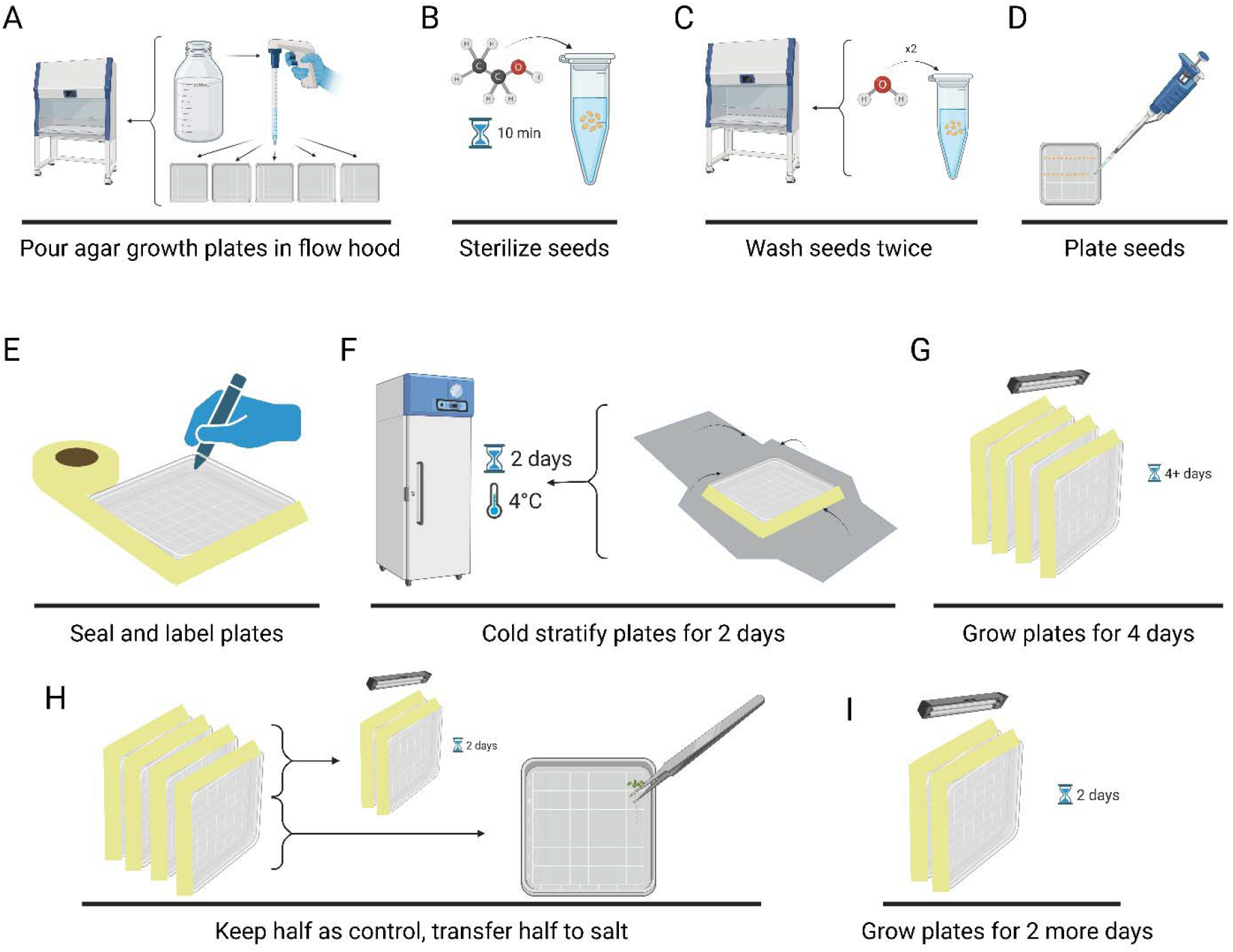
Growth of *A. thaliana* Seedlings. (**A**) In a flow hood, pour 50mL of melted growth media into each plate. (**B**) Put seeds in an Eppendorf tube, add 70% ethanol, mix, and let sit for 10 minutes. (**C**) In the fume hood, remove ethanol and rinse twice with water. (**D**) Use a pipette to plate seeds on hardened agar in two rows of 14 per plate. (**E**) Seal plates with breathable tape and label with a sharpie. (**F**) Wrap plates in aluminum foil and cold stratify in a 4**°**C fridge for 2 days. (**G**) Grow plates upright for 4 days under growth lights. (**H**) Keep half of plates as control plates, transfer the other half onto 100mM NaCl plates in a flow hood, then seal and label. (**I**) Grow plates for 2 more days before imaging.

**Figure 2.**
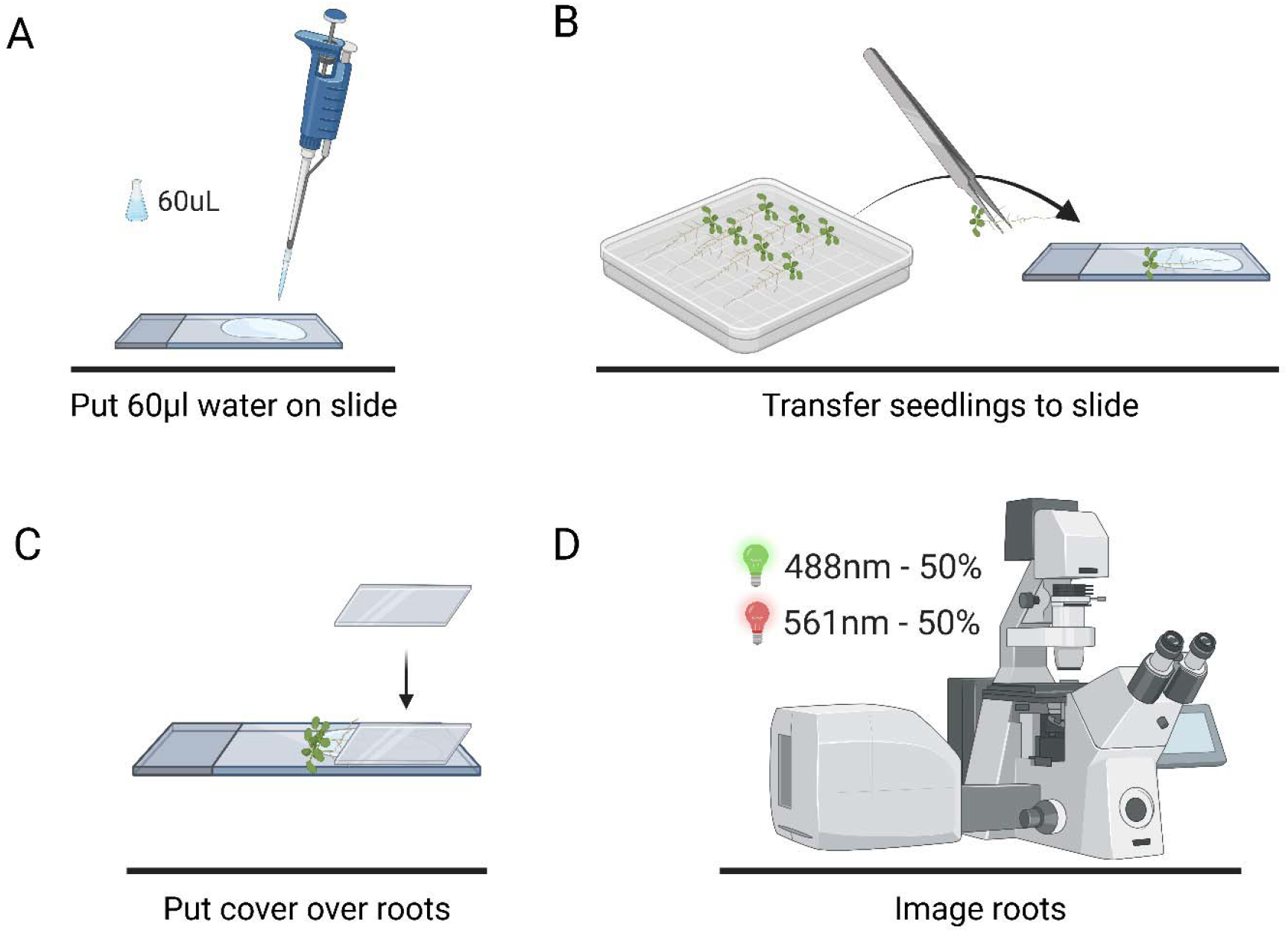
Preparation of seedlings for imaging. (**A**) Use a pipette to put 60μl water onto a slide. (**B**) Use tweezers to transfer seedlings to the slide, placing roots but not leaves into the water. (**C**) Place a 22x22mm slide cover over only the roots. (**D**) Set confocal microscope laser balance to 50% for 488nm and 50% for 561nm, then image roots.

**Figure 3.**
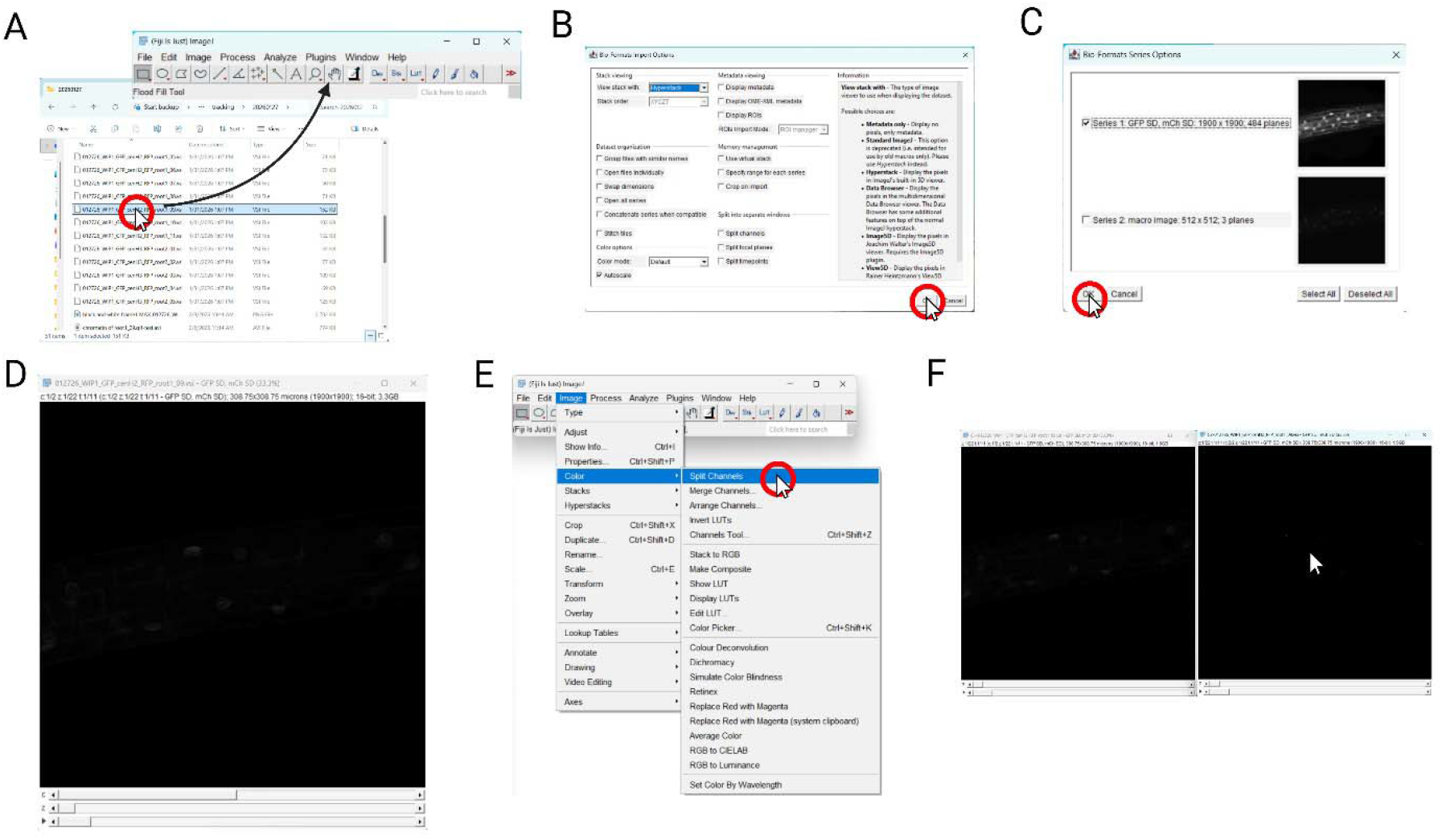
Loading data and splitting channels. (**A**) Drag the .vsi file onto the ImageJ window. (**B**) Click “Ok” on “Bio-Formats Import Options” window. (**C**) Click “Ok” on “Bio-Formats Series Options” window. (**D**) Example of an opened image window. (**E**) Split signals by navigating to image, color, and split. (**F**) The original image will split, click on the one with chromatin.

**Figure 4.**
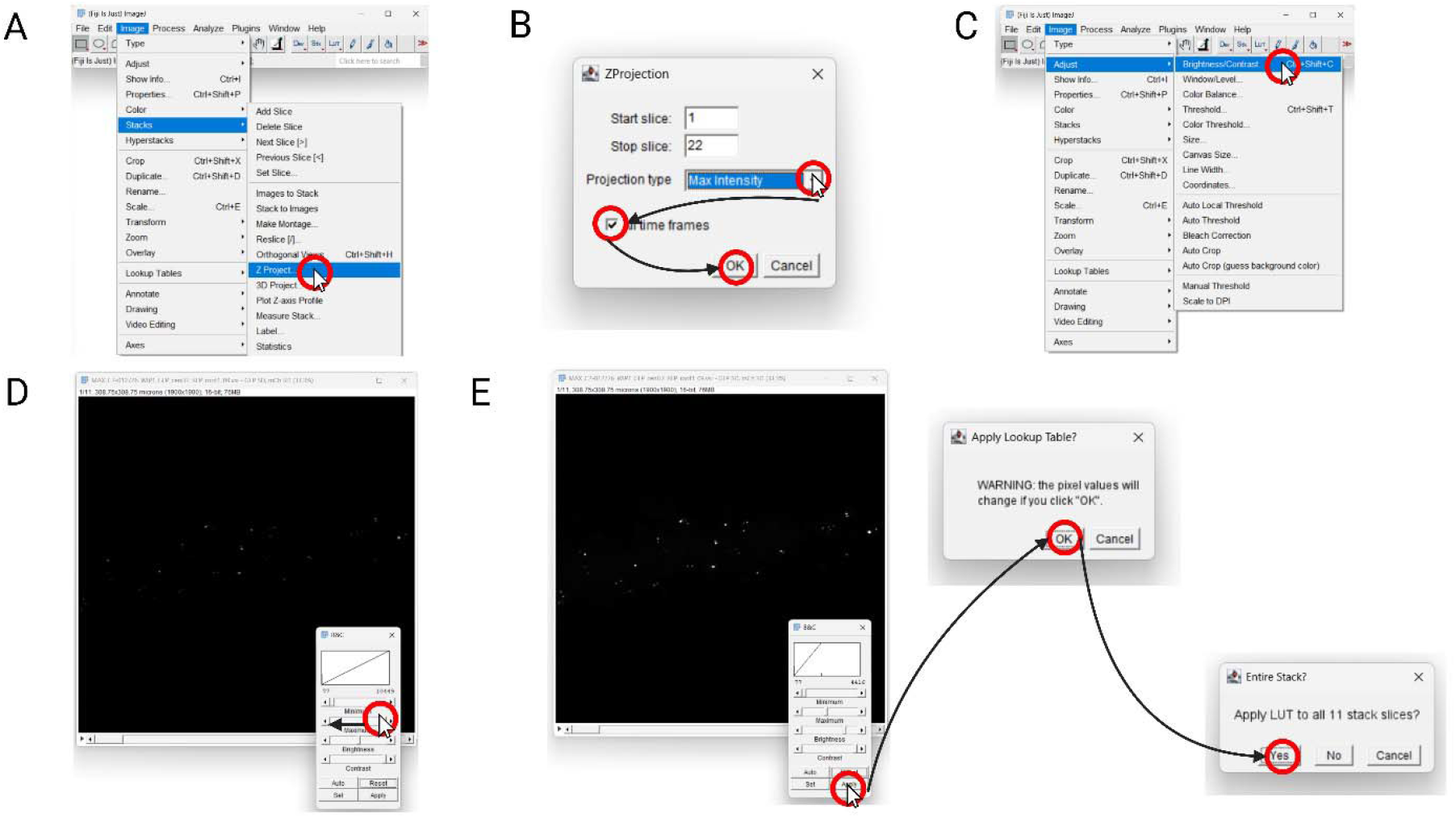
Z-projection and brightness editing. (**A**) Apply Z-project operation by navigating to image, stacks, and “Z project…”. (**B**) Change projection type to “Max Intensity” and ensure “All Time Frames” is active, then click “Ok”. (**C**) Adjust contrast by navigating to image, adjust, and “Brightness/Contrast”. (**D**) Slide the “Maximum” bar to the left until chromatin is easily visible. (**E**) Click “Apply”, “Ok”, then “Yes”.

**Figure 5.**
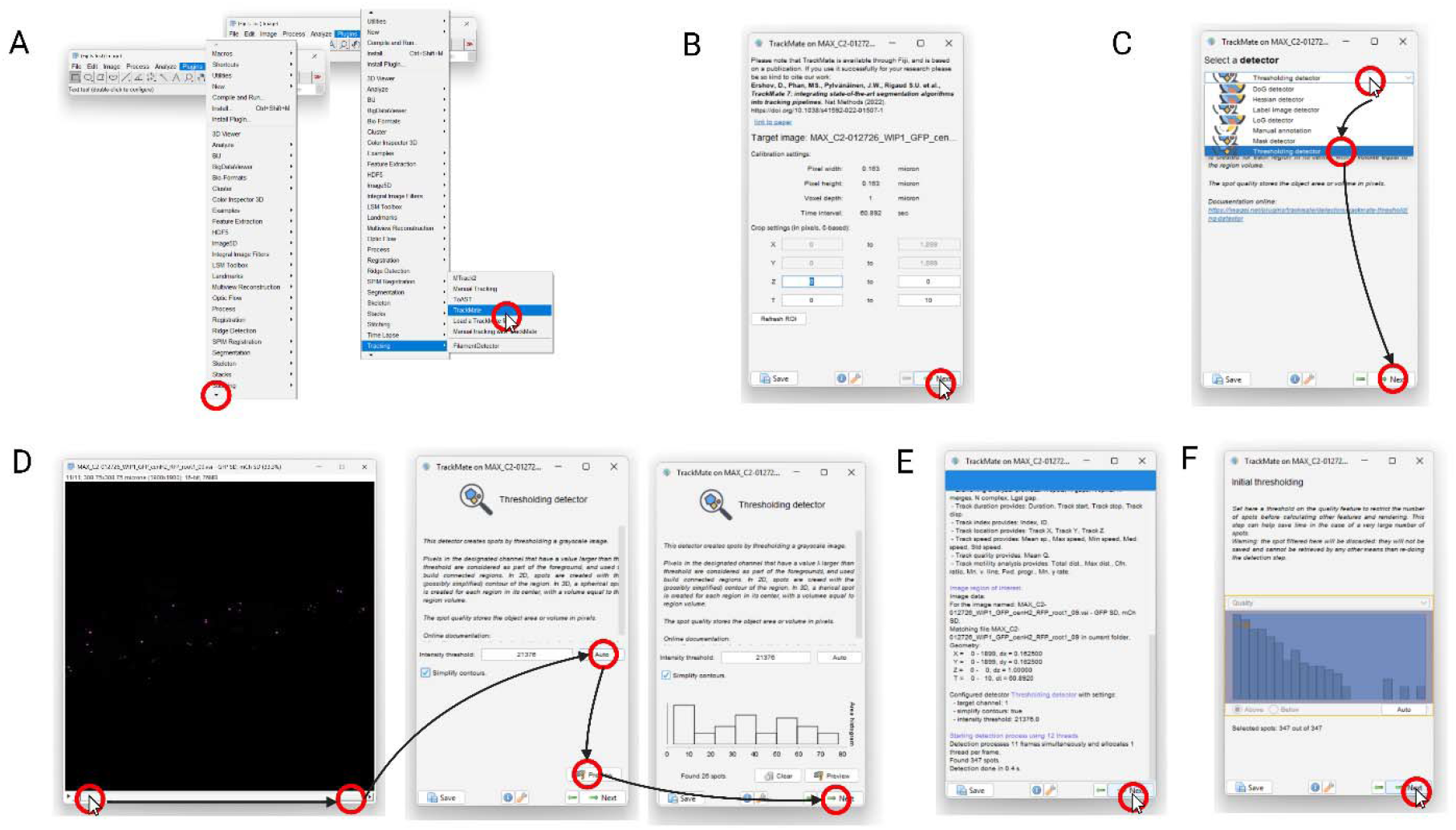
Trackmate thresholding. (**A**) Navigate to plugins and click the down arrow to uncover tracking, then click tracking and “TrackMate”. (**B**) Click “Next”. (**C**) Select “Thresholding Detector” and click “Next”. (**D**) On the chromatin window, drag the time-bar to move to the last frame of the timelapse, then click “Auto” on the TrackMate window, then “Preview”, then “Next”. (**E**) Click “Next”. (**F**) Click “Next”.

**Figure 6.**
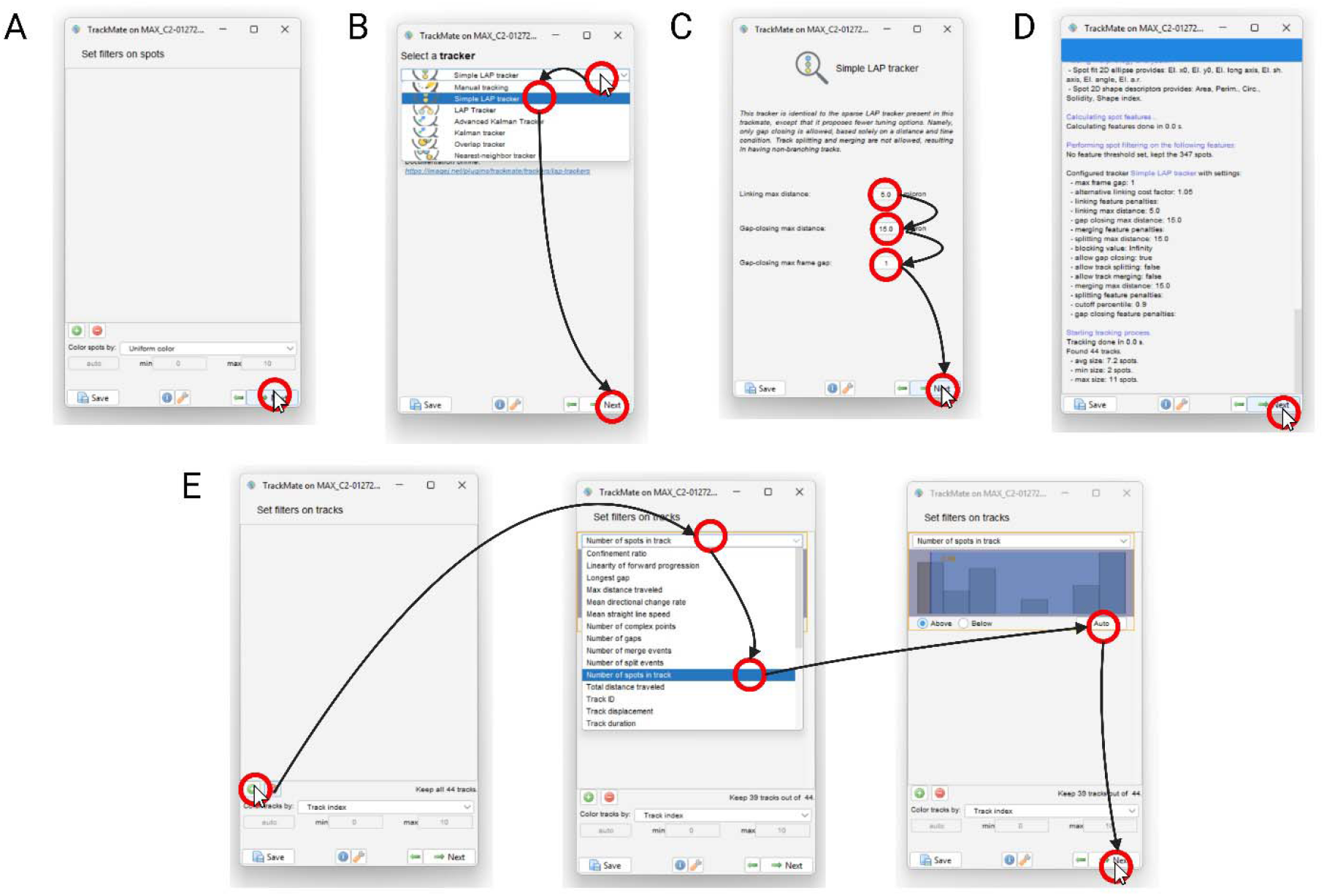
Trackmate tracking. (**A**) Click “Next”. (**B**) Select “Simple LAP tracker” and click “Next”. (**C**) Set linking max distance to 5 microns, gap-closing max distance to 15 microns, and gap-closing max frame gap to 1, then click “Next”. (**D**) Click “Next”. (**E**) Click the green “+” button, then choose the “Number of spots in track” mode and press “Auto”.

**Figure 7.**
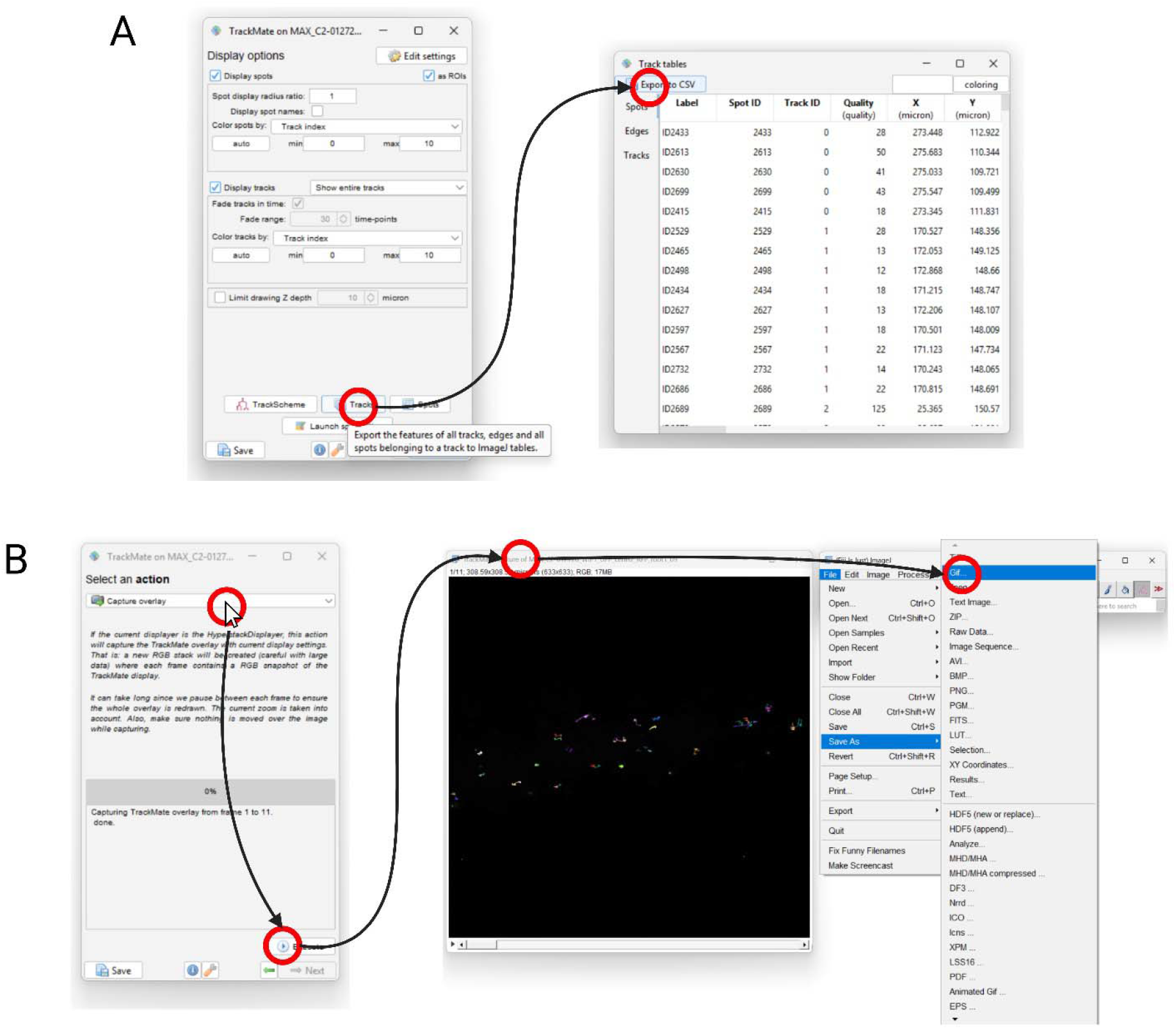
Downloading tracking data. (**A**) Click “Tracks”, then “Export to CSV”. (**B**) Skip the plotting page. Change the mode to “Capture overlay” and click “execute”. Click on the created window, then navigate to file, save as, “GIF…” to save the tracking video.

## Results

### Live cell imaging of nuclear envelope and chromatin dynamics

We generated a dual fluorescent marker line, which includes outer nuclear envelope protein WIP1-GFP and CENTROMERIC HISTONE H3 (CENH3) with mRFP (**Figure 8**) (11,12). 4-days-old seedlings were transferred to the control plate or 100mM NaCl containing plates for 2 days. After 2 days of incubation, there is a clear root growth inhibition observed for salt treatment. Both control and salt-treated roots were used to perform live cell imaging and capture time lapse movies for 5-10 minutes with a minute of interval between acquisition (**Supplemental movies 1 and 2**). In contrast to control, salt treated root cells appear stressed and chromatin morphology is also altered a little bit. Many of these chromatin appear little bigger than control conditions (**Figure 8**). These observations clearly indicate the effect of salt stress in the root at cellular level.

**Figure 8.**
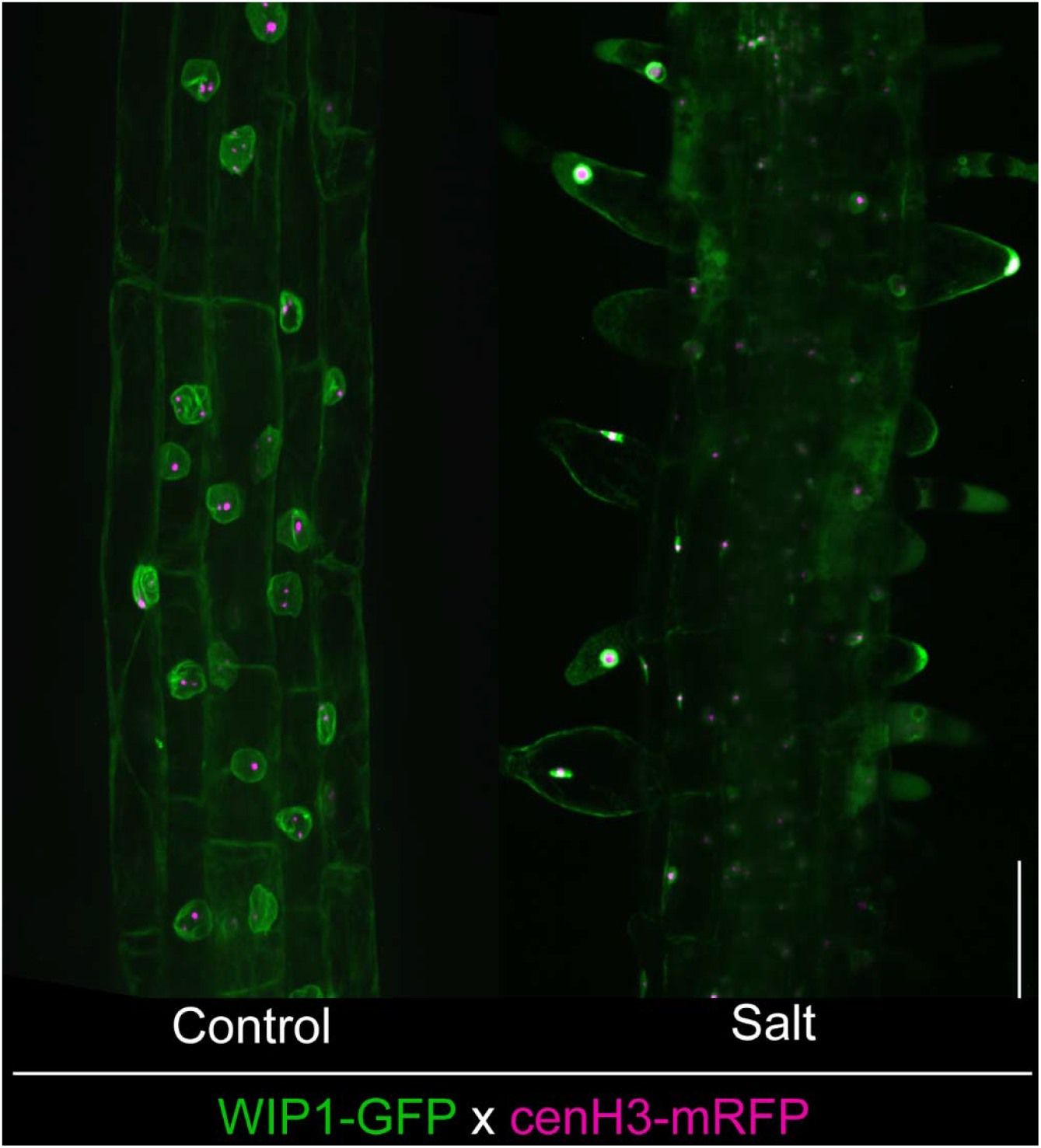
Visualization of nuclear envelope and chromatin. The nuclear envelope and chromatin was visualized using WIP1-GFP and CENH3-mRFP, respectively, in control and salt-treated *A. thaliana* root. Scale bar = 100 µm.

### Quantification of chromatin dynamics from the live cell movies

Acquired movies were processed with ImageJ/Fiji. The first and most important step is to segment the chromatin dot (**Supplemental movies 3, 4, 5, and 6**). The segmented chromatin dots were analyzed using the TrackMate plugin and the displacement between each time point or intervals were measured. This displacement (micron) data was considered over time (sec) to quantify the velocity (micron/sec) for both control and salt-treated conditions. Interestingly, the chromatin velocity differed significantly between control and salt-treated conditions (**Figure 9**). This result provides new insights about chromatin dynamics in *A. thaliana* root and will be useful to test during other biotic and abiotic stress conditions.

**Figure 9.**
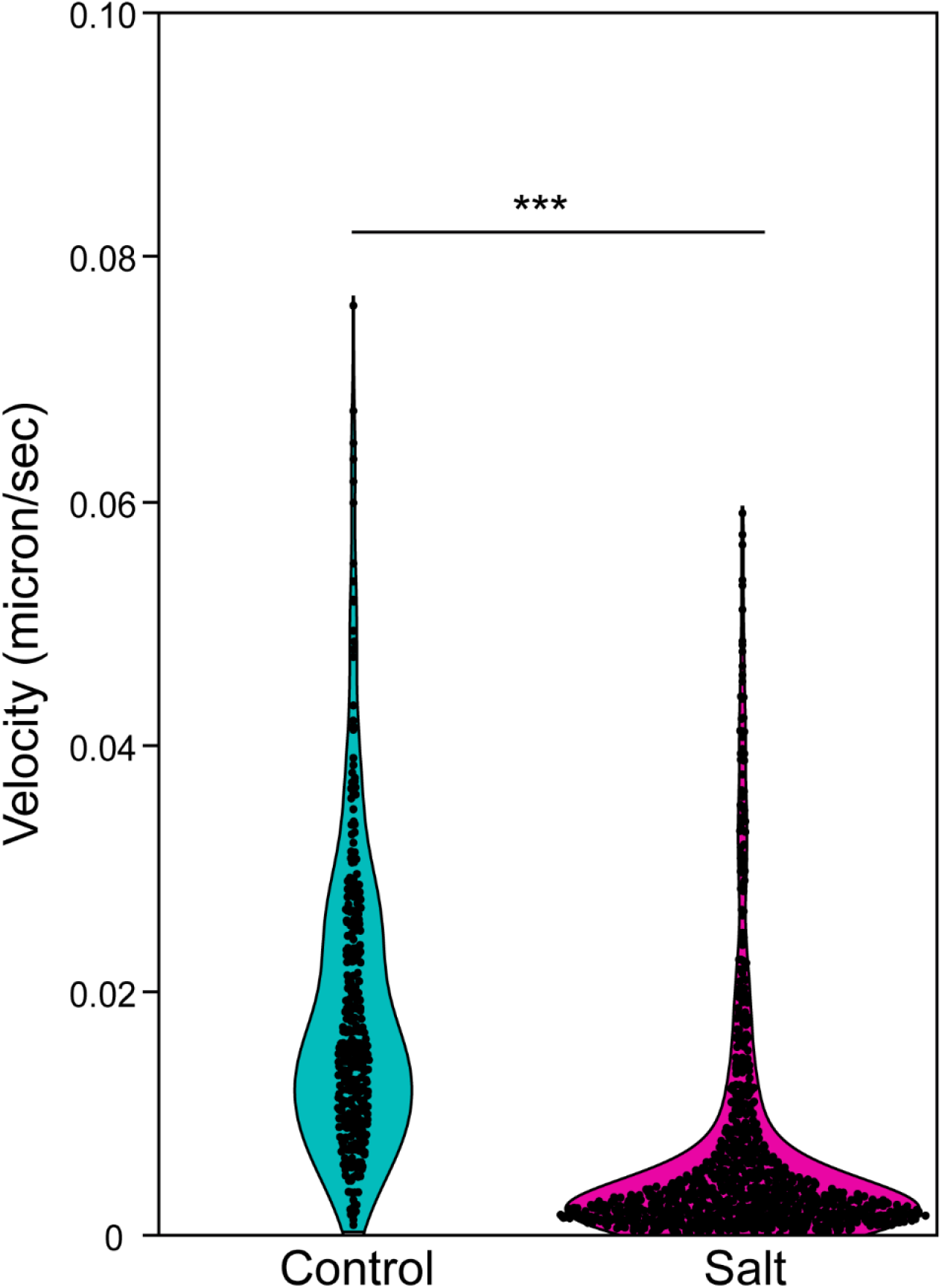
Quantification of chromatin velocity. The segmented chromatin velocity was measured based on the displacement (micron) between two time points over time (sec). The chromatin velocity (micron/sec) is plotted for control and salt-treated roots. Student’s t-test was performed, where asterisks represent the statistical significance between the means for each treatment. ***P < 0.001.

## Discussion

The method presented here is ideal to track chromatin dynamics. As this method relies on live cell imaging markers, it is more reliable than fixed sample and immunostained protocol. As a result, it is most important to have a fluorescent line with consistent signals during the entire process. For the image acquisition, the spinning disk confocal microscope will be ideal in contrast to laser scanning confocal microscope. During the image processing, the segmentation of chromatin dot is another critical step (13). Users need to pay attention whether segmented chromatin dots are segmented throughout all the time points from the time lapse movies. If not, it is very difficult to get a good and reliable dataset from TrackMate.

We have demonstrated the capability of using double fluorescent marker line, live time lapse imaging, and quantification using TrackMate for *A. thaliana* root system. This method will be applicable across tissue and cell type including model and crop plants (14,15). The recent advancement of single cell sequencing data will help to select the cell-type specific expression of chromatin variants (16,17). It will help us to design better cell-type specific fluorescent markers for studying chromatin dynamics. Furthermore, this presented method will be very useful to investigate chromatin dynamics during different environmental stress and stress recovery.

The current method relied on live cell movies captured for 10 minutes in a horizontal confocal stage. As a result, it is challenging to keep the plants alive for a long time. Furthermore, plant roots grow vertically. Here, roots were grown vertically, but imaged horizontally. Using a confocal microscope with the vertical stage will solve this problem and it will be an ideal future endeavor to improve this method. As the root keeps growing, it is difficult to keep the root within a perfect focus during the imaging over a certain period. An automated tracking software to keep nucleus and chromatin on focus during the entire imaging will ensure better output movies. We have also observed that after salt stress, the root cell morphology is altered, and consequently the capturing time lapse movies become harder. This challenge can be circumvented by playing around with a time frame, when stress affects chromatin dynamics, but does not alter the overall cellular morphology. Currently, we cannot categorize the chromatin dynamics based on cell-type specific manners. In the near future, developing cell-type specific double fluorescent marker lines will help to solve this problem and ensure better time lapse movies.

Overall, the method described here will be useful to chromatin dynamics from plant and non-plant organisms across cell-types, developmental stages, stress conditions, and during diseased.

## Acknowledgements

Authors thank Miki Fujita and EunKyoung Lee of UBC Bioimaging Facility (RRID: SCR_021304) for their kind support.

The research at Ashraf lab is funded by the NSERC Discovery grant (RGPIN-2025-04277), Canada Foundation for Innovation (CFI) John R. Evans Leaders Fund (JELF), British Columbia Knowledge Development Fund (BCKDF), and start-up grant provided by the University of British Columbia. Joh Demura-Devore is supported by the Work Learn International Undergraduate Research award by the University of British Columbia.

